# Volatile profiling distinguishes *Streptococcus pyogenes* from other respiratory streptococcal species

**DOI:** 10.1101/2023.04.13.536753

**Authors:** Amalia Z. Berna, Joseph A. Merriman, Leah Mellet, Danealle K. Parchment, Michael G. Caparon, Audrey R. Odom John

**Affiliations:** Department of Pediatrics, Washington University School of Medicine, St. Louis, MO 63110, USA; Department of Pediatrics, Children’s Hospital of Philadelphia, Philadelphia, PA 19104, USA; Department of Molecular Microbiology, Washington University School of Medicine, St. Louis, MO 63110, USA; Microbiome Therapies, Stanford University, Palo Alto, CA 94304, USA; Vaccine Research Center, National Institute of Allergy and Infectious Diseases, Bethesda, MD 20892, USA; Perelman School of Medicine, University of Pennsylvania, Philadelphia, PA 19104, USA

**Author notes:** Correspondence to: Audrey R. Odom John MD PhD (current address), Children’s Hospital of Philadelphia, 3501 Civic Center Blvd, CTRB 10100, Philadelphia PA 19104-4318.

**Keywords:** pharyngitis, volatile organic compounds, Group A streptococci, *Streptococcus pyogenes*

## Abstract

Sore throat is one of the most common complaints encountered in the ambulatory clinical setting. Rapid, culture-independent diagnostic techniques that do not rely on pharyngeal swabs would be highly valuable as a point-of-care strategy to guide outpatient antibiotic treatment. Despite the promise of this approach, efforts to detect volatiles during oropharyngeal infection have yet been limited. In our research study, we sought to evaluate for specific bacterial volatile organic compounds (VOC) biomarkers in isolated cultures *in vitro*, in order to establish proof-of-concept prior to initial clinical studies of breath biomarkers. A particular challenge for diagnosis of pharyngitis due to *Streptococcus pyogenes* is the likelihood that many metabolites may be shared by *S. pyogenes* and other related oropharyngeal colonizing bacterial species. Therefore, we evaluated whether sufficient metabolic differences are present that distinguish the volatile metabolome of Group A streptococci from other streptococcal species that also colonize the respiratory mucosa, such as *S. pneumoniae* and *S. intermedius*. In this work, we identify candidate biomarkers that distinguish *S. pyogenes* from other species, and establish highly produced VOCs that indicate presence of *S. pyogenes in vitro*, supporting future breath-based diagnostic testing for streptococcal pharyngitis.

**IMPORTANCE:** Acute pharyngitis accounts for approximately 15 million ambulatory care visits in the USA. The most common and important bacterial cause of pharyngitis is *Streptococcus pyogenesis*, accounting for 15% to 30% of pediatric pharyngitis. Distinguishing between bacterial and viral pharyngitis is key to management in US practice. Culture of a specimen obtained by throat swab is the standard laboratory procedure for the microbiologic confirmation of pharyngitis, however this method is time consuming which delays appropriate treatment. If left untreated, *S. pyogenes* pharyngitis may lead to local and distant complications. In this study, we characterized the volatile metabolomes of *S. pyogenes* and other related oropharyngeal colonizing bacterial species. We identify candidate biomarkers that distinguish *S. pyogenes* from other species and provides evidence to support future breath-based diagnostic testing for streptococcal pharyngitis.

## INTRODUCTION

Acute pharyngitis is a common outpatient medical condition that leads to approximately 15 million healthcare visits per year in the United States (1). Infection with *Streptococcus pyogenes* (group A beta-hemolytic streptococcus) is the most common bacterial cause of acute pharyngitis and is responsible for up to 15% of cases among adults and 30% of cases among children (1). However, a major challenge in outpatient management of acute pharyngitis is the inability to easily distinguish between bacterial and viral etiologies on the basis of clinical presentation alone. Oral treatment with a beta-lactam antibiotic, specifically penicillin or amoxicillin, is recommended for tonsillopharyngitis due *S. pyogenes*, in order to reduce rates of post-infectious acute rheumatic fever.

Standard clinical practice to evaluate for streptococcal pharyngitis depends on direct provider sampling of the posterior oropharynx (1), followed by point-of-care rapid testing for streptococcal antigens as well as bacterial culture. While antigen detection together with culture identify *S. pyogenes* pharyngitis with a high sensitivity, inadequate oropharyngeal sampling and cross-reactivity with other oral colonizers can lead to false negative or false positive results. Inappropriate antibiotic use, particularly for viral respiratory infections, is a key factor in rising rates of antimicrobial resistance worldwide (2, 3). The Centers for Disease Control and Prevention estimates that over one-third of all outpatient antibiotics prescribed to 0-19 year old for respiratory symptoms, including pharyngitis, may be unnecessary (4).

One possible alternative approach to standard microbiological diagnosis takes advantage of the distinct metabolic capabilities of different bacterial species. As bacteria utilize different substrates for growth, by-products of these utilized substrates may yield additional chemical biomarkers, such as volatile organic compounds (VOCs), that allow for identification, classification, and discrimination of microorganisms (5–7). Many bacteria produce characteristic volatile metabolites that can be detected through chemical analysis of the headspace gas above bacterial cultures. These VOCs contribute to the characteristic odor profiles already well associated with particular bacterial species. For example, the canonical grape-like odor associated with *Pseudomonas aeruginosa* is caused by production of 2-aminoacetophenone, whereas the distinct odor associated with cultured *E. coli* is that of indole. Although the biological functions of most bacterial VOCs are yet unexplored, recent evidence suggests they may function in interbacterial communication, defense, and growth promotion (6). Highly sensitive molecular detection based on gas chromatography-mass spectrometry (GC-MS) instruments allow for unbiased identification of organism-specific VOC biomarkers (8). Many human bacterial pathogens can be cultured in liquid or solid medium, enabling straightforward VOC detection from axenic culture before investigating a more complex *in vivo* system. This advance in technology and abundance of information holds the promise for diagnosing infections *in situ* using volatiles—for example, directly from an infected wound, a urine sample, or the breath (9–11).

Volatile analysis of human breath has already been successful in pilot studies for the detection of infections such as malaria and pulmonary tuberculosis (9, 10, 12) and is commercially available for the detection of *Helicobacter pylori* infection (13), demonstrating the validity and widespread applicability of this diagnostic technique. When evaluating for streptococcal pharyngitis, oropharyngeal volatiles are likely to reflect the metabolic activity of the pathogen, at least in part, and may also comprise any host-associated volatiles triggered as a result of infection. However, interpretation of oropharyngeal volatile composition in a clinical setting may be challenging, due to the complex oropharyngeal microbial community found in the oropharynx and other endogenous and exogenous sources of VOCs.

In this study, we sought to evaluate for specific bacterial VOC biomarkers in isolated cultures *in vitro*, in order to establish proof-of-concept prior to initial clinical studies of breath biomarkers. A particular challenge for diagnosis of *S. pyogenes* is the likelihood that many metabolites may be shared by *S. pyogenes* and other related oropharyngeal colonizing bacterial species. For this reason, we performed *in vitro,* culture-based, examination of the volatile profile of cultured *S. pyogenes*, compared with two other common respiratory streptococcal species, *S. intermedius* and *S. pneumoniae*. In this work, we identify candidate biomarkers that distinguish *S. pyogenes* from other species, and establish highly produced VOCs that indicate presence of *S. pyogenes in vitro*, supporting future breath-based diagnostic testing for streptococcal pharyngitis.

## MATERIALS AND METHODS

### Isolate collection

*Streptococcus pyogenes* strains were supplied by the culture collection of Dr. Michael Caparon. Strains chosen for this study were isolated from pediatric patients with streptococcal pharyngitis, treated at Saint Louis Children’s Hospital between 2007 and 2012 as part of an unpublished work. *Streptococcus pneumoniae* (Spneu1) was donated by Dr. Malcolm Winkler, the isolate originated from the American Type Culture Collection (ATCC BAA-255), while Spneu2 and Spneu3 originated from nasopharyngeal swabs collected at Saint Louis Children’s Hospital, graciously provided by Dr Celeste Morley (14) (Table 1). Samples were collected at Saint Louis Children’s Hospital under approval by the Washington University School of Medicine Institutional Review Board/Human Research Protection Office (14). *Streptococcus intermedius* isolates were acquired from ATCC (Table 1). Species identification was performed using standard clinical microbiological testing.

**Table 1.** Bacterial species and strains used in this study.

| Species | Sample ID | Source | Reference |
| --- | --- | --- | --- |
| *Streptococcus pyogenes* | Spyo1 | Pharyngitis isolate | This study |
| *Streptococcus pyogenes* | Spyo2 | Pharyngitis isolate | This study |
| *Streptococcus pyogenes* | Spyo3 | Pharyngitis isolate | This study |
| *Streptococcus intermedius* | Sint1 | ATCC31412 | This study |
| *Streptococcus intermedius* | Sint2 | ATCC9895 | This study |
| *Streptococcus intermedius* | Sint3 | ATCC27335 | This study |
| *Streptococcus pneumoniae* | Spneu1 | Originally labeled EL59 by Malcolm Winkler | (37) |
| *Streptococcus pneumoniae* | Spneu2 | Nasopharyngeal isolate | (14) |
| *Streptococcus pneumoniae* | Spneu3 | Nasopharyngeal isolate | (14) |

### Culture conditions and sample preparation

Low passage-number isolates were stored at −80 °C in 25% glycerol until the time of analysis. Two days before headspace sampling, isolates were plated directly from −80°C stock on Todd Hewitt plus yeast extract (THY) solid agar plates. Strains were grown under anaerobic conditions using BD GasPak™ EZ system (Becton, Dickinson and Company Sparks, MD, United States) at 37°C, overnight. Isolates were then inoculated into 10mL THY broth in a 20 mL air-tight glass headspace vial and sealed with silicone/PTFE screw caps (Sigma-Aldrich, St Louis, MO, United States). Samples were incubated in stationary culture at 37°C for 24 hours, after which headspace collections were made. Three independent clinical isolates were evaluated for each bacterial species. For each bacterial isolate, three independent experimental repeats on three separate days were prepared and the headspace analyzed as described below, along with an uninfected media control collected on each day of analysis.

### Concentration and analysis of volatile compounds

Volatile metabolites were concentrated using headspace solid-phase micro extraction (HS-SPME) and separated and analyzed via gas chromatography-mass spectrometry (GC-MS) (Figure 1). Headspace extraction was carried out for 30 min with a HS-SPME fiber composed of fused silica partially cross-linked with 50/30 µm Divinylbenzene/Carboxen/Polydimethylsiloxane (DVB/CAR/PDMS). After absorption, headspace volatiles were transferred to the GC injection port, which was equipped with a 0.8 mm i.d. splitless glass liner, at 250°C. Desorbed volatile compounds were separated in an Agilent 7890A GC, equipped with a DB-5MS (Agilent Technologies Inc., California U.S.) fused silica capillary column (30 m × 0.25 mm, 0.25 μm film thickness). The oven temperature was programmed to rise from 35°C (held for 4 min) to 65°C at 2.5°C/min and rise again to 100°C at 5°C/min and finally to 230°C at 30°C/min (held for 2 min). The GC column output was fed into an Agilent 5975C mass selective detector. The GC-MS transfer line was heated at 300°C with He as the carrier gas (1 mL/min). Mass spectrometry was performed in electron impact mode at 70 eV scanning over the range m/z = 35-350 Da.

**Figure 1:**
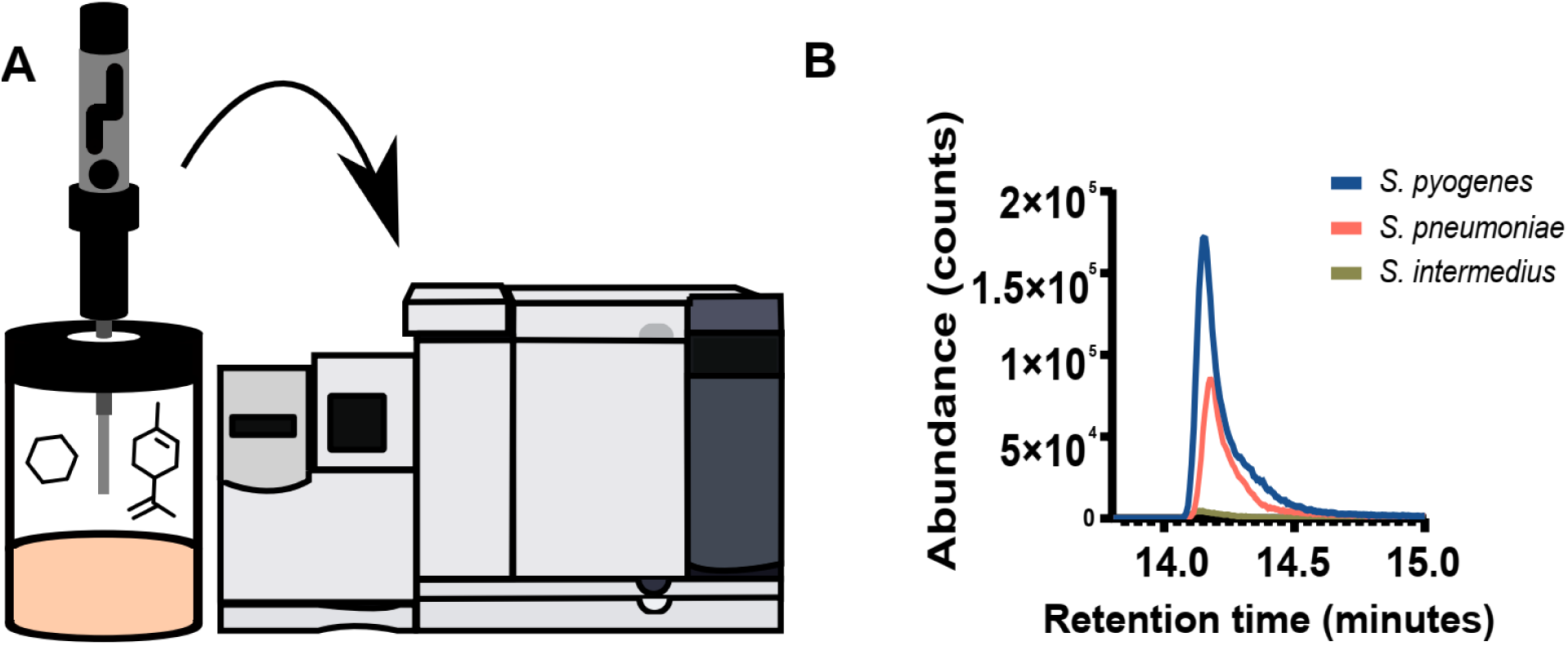
(A) Scheme of volatile organic compounds collection from glass vials using headspace solid phase micro-extraction and subsequent analysis with GC-MS. (B) Abundance of characteristic biomarker, benzaldehyde, found in *S. pyogenes, S. intermedius,* and *S. pneumoniae.* Benzaldehyde by the base ion peak (m/z 106) area is shown.

To correct for day-of-analysis effects on GC-MS data (15), an external standard EPA 8240B Calibration Mix Sigma-Aldrich (USA) was used each day of GC-MS analysis to normalize for day-specific effects. A 20 μg/mL EPA calibration solution was made up in methanol (HPLC grade) on each day of measurement. 1 mL of this solution was placed in a 20 mL air-tight glass headspace vial and sealed with silicone/PTFE screw cap (Sigma-Aldrich, St Louis, MO, United States) and run at the start of every GC-MS sequence.

### Compound reporting

Chromatographic peaks were assigned putative identifications based on mass spectral matching. Specifically, putative compound identifications were assigned to peaks with a forward match score of ≥600/1000 relative to the National Institute of Standards and Technology (NIST) 2011 mass spectral library (16).

### Statistical Analyses

Data were analyzed by using interactive XCMS Online (17), which is freely available at https://xcmsonline.scripps.edu. Metabolite features were defined as ions with unique m/z and retention-time values. For XCMS processing of GC-MS data, parameter settings were as follows: centWave for feature detection (Δ m/z = 100 ppm, minimum peak width = 5 s, maximum peak width = 10 s, and signal/noise threshold= 6); obiwarp settings for retention-time correction (profStep = 1); and parameters for chromatogram alignment, including mzwid = 0.25, minfrac = 0.5, and allowable retention time deviationsbw = 10. The relative quantification of metabolite features was based on EIC (extracted ion chromatogram) areas. In total 1420 features were detected.

We applied multiple group analysis to compare the means of multiple independent groups (in our case, 3 groups) and enable the identification of metabolite features whose variation pattern is statistically significant. To evaluate the metabolite variation across different experimental groups, we used the nonparametric alternative, Kruskal– Wallis test (p-value threshold 0.05). Principal Component Analysis (PCA) was used to visualize variance among samples using the molecules detected from Kruskal–Wallis test. For PCA, features with m/z >150 a.m.u., retention time >26 min and p>0.5 were exclude from this analysis. Large masses and compounds that leave the chromatographic column after 26 min are mainly contaminants. Additionally, features with an area <1000 counts were considered at or below the limit of detection and were not considered for analysis. Each feature was then normalized to the 2-hexanone external standard. A feature in a bacteria group was retained if data was available in more than 50% of the samples. This filtering process lead to 418 features for PCA. PCA was performed with Matlab (version 8.0). Hierarchical clustering of the discriminatory volatiles was carried out using ClustVis as previously described (18).

## RESULTS

We sought to evaluate whether sufficient metabolic differences are present that distinguish the volatile metabolome of Group A streptococci from other streptococcal species that also colonize the respiratory mucosa, such as *S. pneumoniae* and *S. intermedius*. To identify candidate species-specific biomarkers of *S. pyogenes*, we performed comparative profiling of volatile organic compounds (VOCs) derived from axenic cultures of clinical isolates of *S. pyogenes, S. pneumoniae*, and *S. intermedius*.

Headspace VOCs were absorbed via solid-phase microextraction, prior to analysis via gas chromatography mass spectrometry (GC-MS) (see scheme, Figure 1A). Successful VOC sampling was confirmed through detection of the canonical bacterial volatile, benzaldehyde, which is readily detected in the headspace of most bacterial cultures (19, 20). Benzaldehyde was detected in headspace sampling from each sample (Figure 1B). Independent analysis of three different clinical isolates for each species, in triplicate, was performed to control for potential strain-specific findings.

Principal components analysis (PCA) was performed using features detected in all samples after Kruskal-Wallis test (n= 418 features), revealing a distinct volatile metabolite signature of *S. pyogenes*, compared to *S. pneumoniae* and *S. intermedius* (Figure 2). These results strongly suggest that each species produces a highly unique metabolic profile that may ultimately be harnessed for diagnosis. To identify distinctive candidate biomarkers for each species, we sought to identify the specific VOCs that significantly contribute to the differences observed by PCA. In total, we identified 27 discriminatory VOCs (q values <0.05), comprised of aldehydes, alcohols, nitrogen-containing compounds, hydrocarbons, ketones, aromatic compounds, esters, ethers, and carboxylic acid (Table 2).

**Figure 2:**
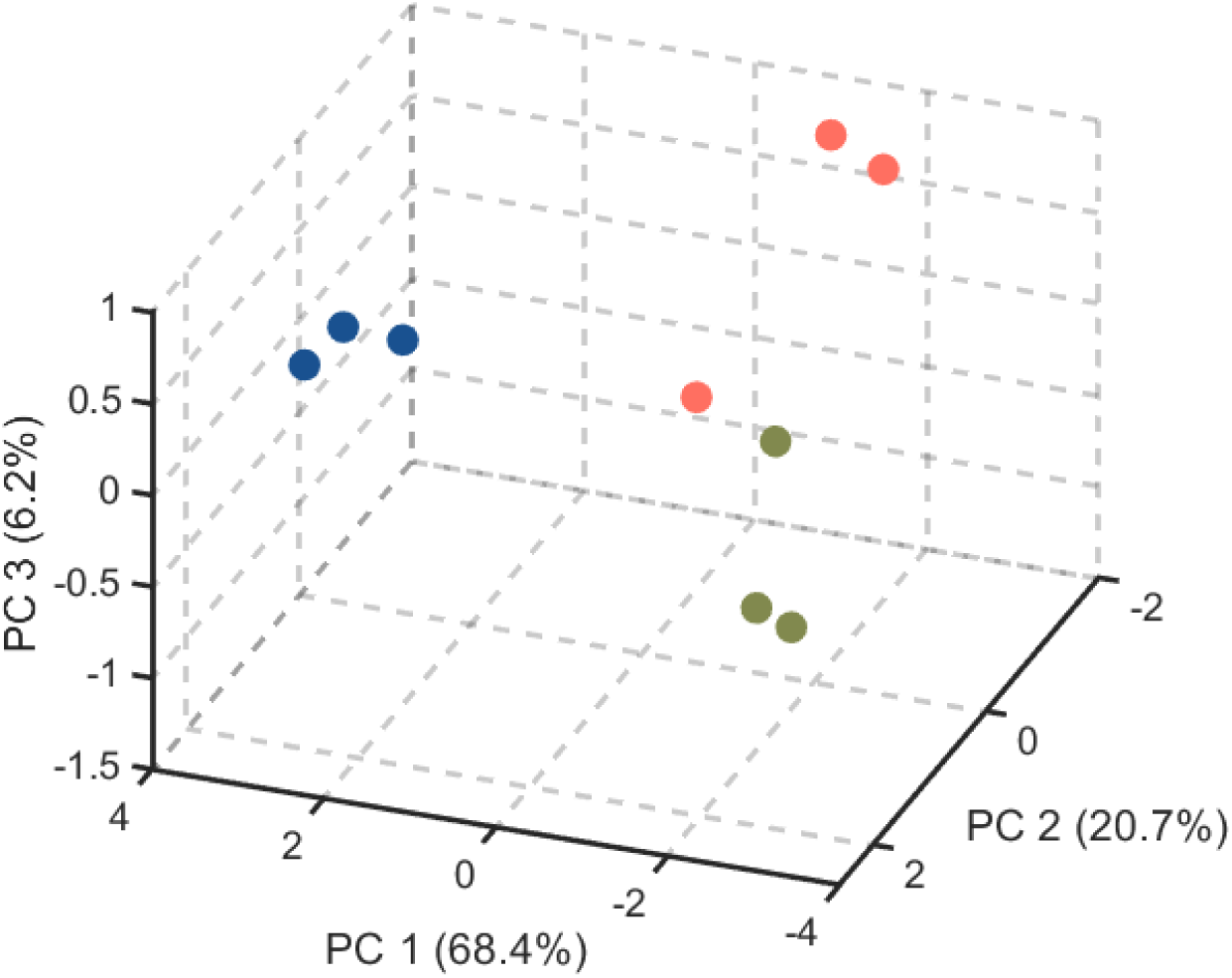
Principal component (PC) score plot for 3 bacterial pathogen groups: S. pyogenes (blue), S. intermedius (green), and S. pneumoniae (coral). Each point in the PCA derives from the mean peak intensities from three biological replicates for each strain.

**Table 2.** Discriminatory volatile organic metabolites found in S. pyogenes, S. intermedius and S. pneumonia cultures.

| VOC number | Rt<sup>†</sup> (min) | Putative name | Ion* (m/z) | Match** | q value |
| --- | --- | --- | --- | --- | --- |
| Aldehyde |  |  |  |  |  |
| 1 | 1.65 | Propanal, 2-methyl | 41 | 769 | 0.0032 |
| 2 | 2.40 | **Butanal, 3-methyl-** | 44 | 938 | 0.0012 |
| 3 | 20.17 | Nonanal | 56 | 762 | 0.0021 |
| Alcohol |  |  |  |  |  |
| 4 | 2.51 | 1-Butanol | 56 | 880 | 0.0034 |
| 5 | 3.87 | 1-Butanol, 3-methyl- | 55 | 926 | 0.0125 |
| 6 | 6.89 | **3-Butyn-1-ol** | 52 | 701 | 0.0016 |
| Nitrogen-containing |  |  |  |  |  |
| 7 | 4.94 | 1, 2-Ethanediamine, N-(2-aminoethyl)- | 73 | 678 | 0.0019 |
| 8 | 3.52 | Ethenamine, N-methylene- | 54 | 855 | 0.0109 |
| 9 | 11.22 | **Pyrazine, 2,5-dimethyl-** | 108 | 912 | 0.0012 |
| 10 | 16.49 | Pyrazine, 2-ethyl-6methyl- | 121 | 802 | 0.0140 |
| 11 | 19.19 | Pyrazine, 2,5-diethyl- | 135 | 747 | 0.0048 |
| 12 | 20.17 | Pyrazine, 3-ethyl-2,5-dimethyl- | 136 | 915 | 0.0015 |
| 13 | 20.16 | 2H-Imidazo[4,5-b]pyrindin-2-one, 1,3-dihydro- | 64 | 853 | 0.0015 |
| Hydrocarbon |  |  |  |  |  |
| 14 | 2.51 | 1, 5-Hexadien-3-yne | 74 | 868 | 0.0216 |
| 15 | 7.5 | **2,4-Dimethyl-1-heptene** | 70 | 897 | 0.0012 |
| Ketone |  |  |  |  |  |
| 16 | 1.4 | Acetone | 43 | 923 | 0.0034 |
| 17 | 1.85 | 2-Butanone | 43 | 871 | 0.0083 |
| 18 | 6.61 | **2,3-Pentanedione** | 43 | 792 | 0.0018 |
| 19 | 20.89 | 2-Nonanone | 41 | 832 | 0.0284 |
| Aromatic |  |  |  |  |  |
| 20 | 4.6 | Toluene | 91 | 806 | 0.0033 |
| 21 | 8.43 | Ethylbenzene | 91 | 875 | 0.0027 |
| 22 | 14.01 | **Benzaldehyde** | 106 | 687 | 0.0015 |
| Ester |  |  |  |  |  |
| 23 | 6.60 | **Butyl acetate** | 41 | 900 | 0.0022 |
| Ether |  |  |  |  |  |
| 24 | 2.03 | **Propane, 2-ethoxy-2-methyl-** | 59 | 868 | 0.0012 |
| 25 | 3.13 | Furan, 2,5-dimethyl | 96 | 897 | 0.0260 |
| Carboxylic acid |  |  |  |  |  |
| 26 | 2.4 | **Acetic acid** | 60 | 937 | 0.0012 |
| Unknown |  |  |  |  |  |
| 27 | 13.27 | Unknown | 41 | - | 0.0015 |

<sup>†</sup>Rt= retention time. \*Ion used for data analysis (m/z). \*\* Peaks were assigned putative identifications based on NIST mass spectral matching. In bold are the VOCs in each chemical family with the lowest q value and their intensities for each bacterial species are plotted in box plots (Figure 4).

To visually represent the relationships between these discriminatory metabolites, we performed hierarchical clustering of metabolites that were identified as significantly different with respect to bacterial species. This analysis yielded two prominent clusters, as shown in Figure 3. The first cluster is characterized by compounds consistently increased in abundance in the headspace gas of *S. pyogenes* cultures, compared to those of *S. pneumoniae* and *S. intermedius*. The second cluster is characterized by compounds consistently decreased in abundance in the headspace gas of *S. pyogenes* isolates. Of note, excellent agreement in volatile fingerprint was observed across all three clinical isolates of *S. pyogenes* evaluated in this study, while strain-dependent volatile production was noted for *S. pneumoniae* and *S. intermedius* (e.g., compare Spneu1 to Spneu2 and Spneu3). However, a larger number of strains will be required to establish the significance of this finding.

**Figure 3:**
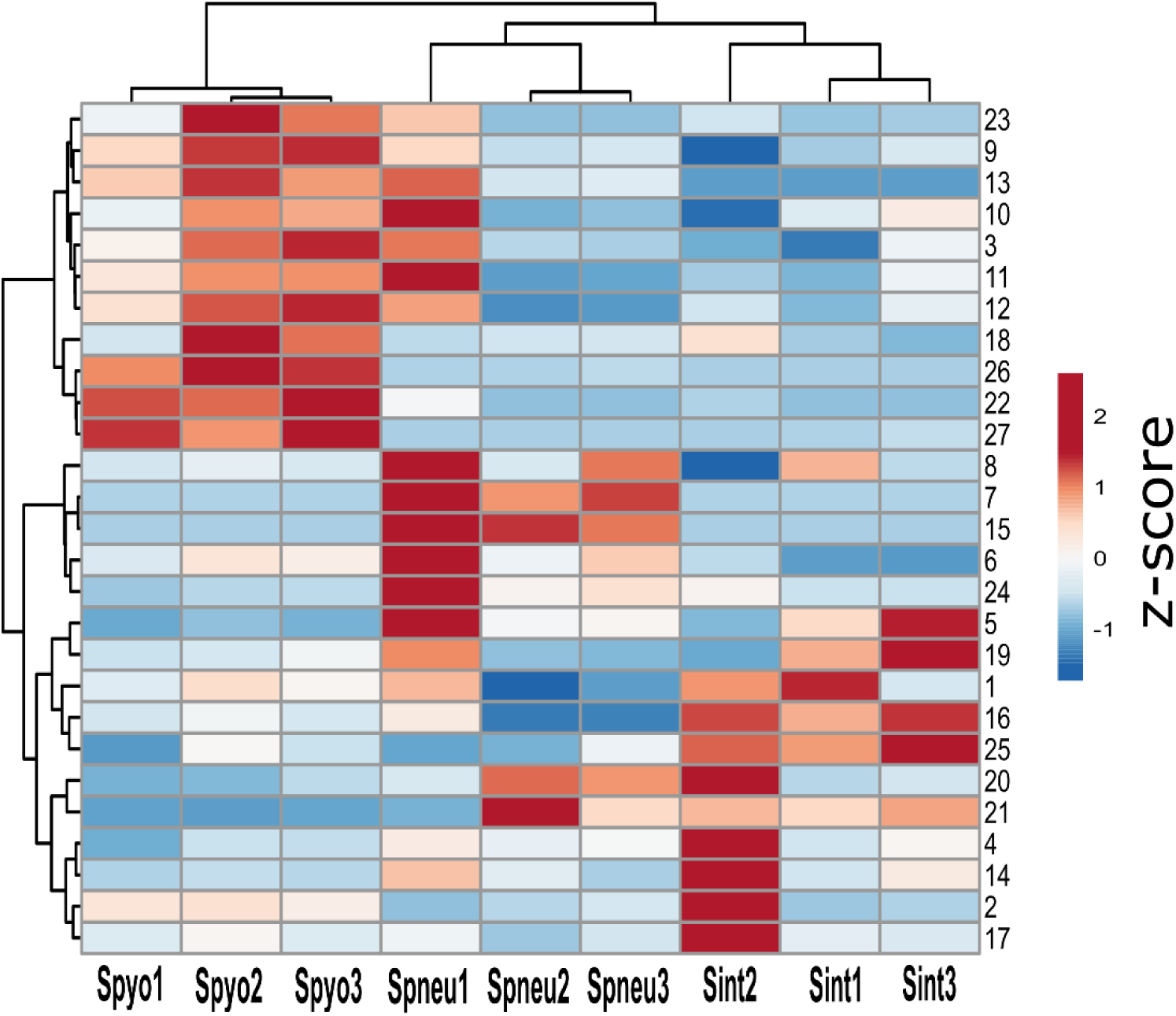
Hierarchical clustering of Z-scores of metabolites significantly different with respect to bacterial species, as calculated from the mean peak intensities from three biological replicates for each strain, normalized to external standard. Volatile numbers on the hierarchical clustering corresponds to discriminant volatiles, as listed in table 2.

Nitrogen-containing compounds represent the largest group of discriminant volatile organic compounds detected (Table 2). Generally, nitrogen-containing compounds (5 of 7 discriminant nitrogen-containing VOCs) were more abundant in the headspace gas of *S. pyogenes* samples (Figure 3). Notably, levels of 2, 5-dimethyl pyrazine were nearly 2.5-fold higher in the headspace gas of *S. pyogenes* compared to that of the other two bacteria species (Figure 4). Three aldehydes were also significantly elevated in the headspace gas of *S pyogenes* (p<0.05) compared to other species of streptococci. In particular, nonanal was higher in *S. pyogenes* compared to the other two species (Figure 3), while 3-methyl-1-butanol was found at significant lower concentrations in *S. pyogenes* in comparison analysis.

**Figure 4:**
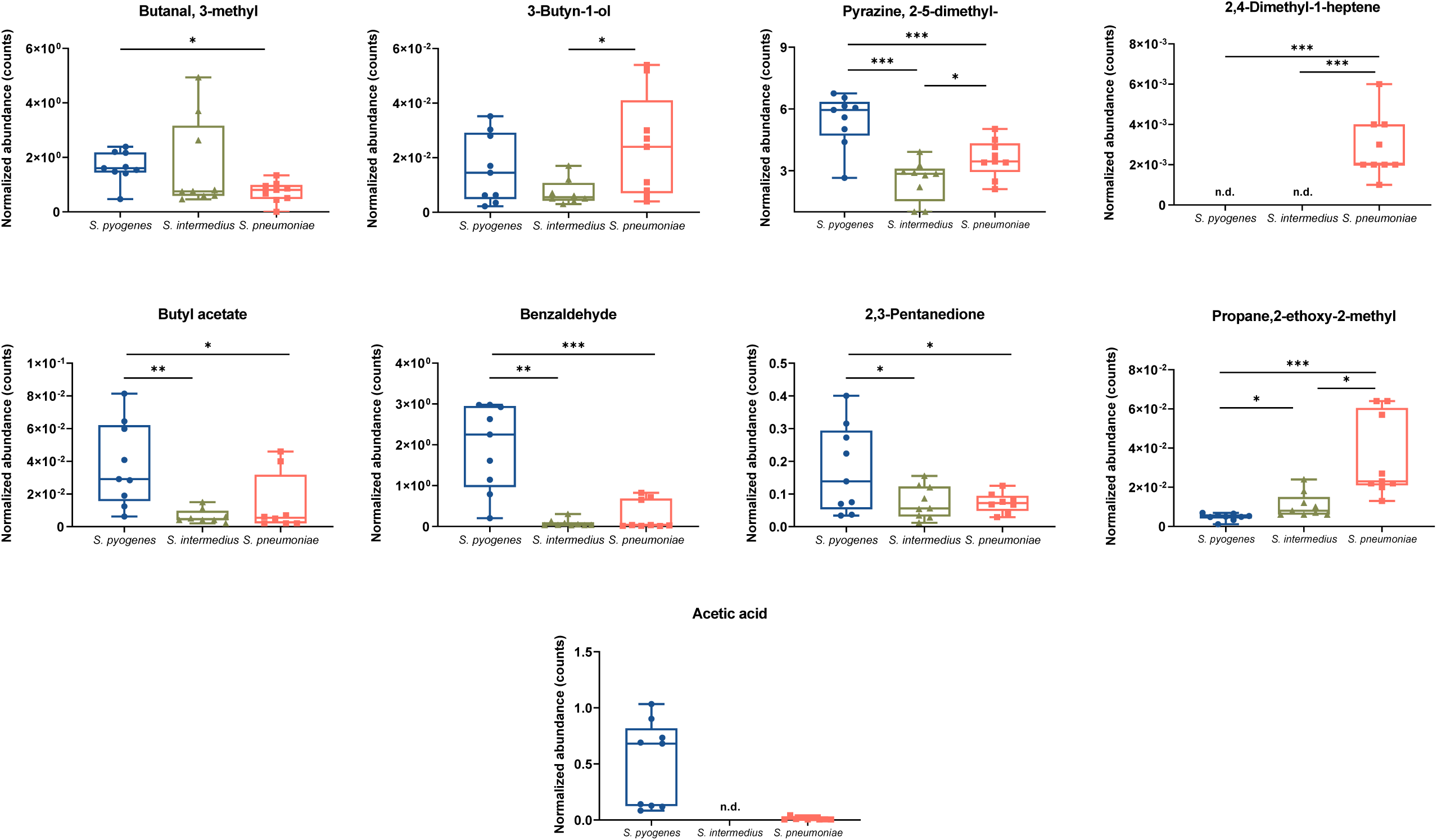
Volatile biomarkers of S. pyogenes cultures. Box plots and Kruskal–Wallis test results for VOCs shown in bold in Table 2. The central mark of each box corresponds to the median, the edges of the box are the 25th and 75th percentiles, the whiskers extend to the most extreme data points not considered outliers. * = P values ≤0.05 (Kruskal–Wallis test) between pairs of bacteria, ** = p values ≤0.01, *** = p values ≤0.001, nd= not detected.

Distinct volatile compounds also characterized the headspace of cultured *S. pneumoniae* and *S. intermedius*. *S. pneumoniae* released a unique volatile organic compound, namely N-(2-aminoethyl)-1, 2-ethanediamine, which was below the limit-of-detection in all samples from *S. pyogenes* and *S. intermedius*. These findings suggest N-(2-aminoethyl)-1, 2-ethanediamine may represent a specific biomarker of *S. pneumoniae* colonization or infection. While robust ketone production was observed across all three streptococcal species, 2,3-pentanedione was markedly increased (2.3 fold) in *S. pyogenes* (Figure 4). Interestingly, although both *S. pyogenes* and *S. intermedius* excrete acetic acid as a result of reducing pyruvate during fermentation (21), *S. intermedius* lacked detectable levels of acetic acid, while this VOC was highly produced by *S. pyogenes* (Figure 4).

Butyl acetate was the only ester that discriminated between the three streptococcal species (Figure 4). Produced by all three species, levels of butyl acetate were substantially (up to 6-hold) more abundant in *S. pyogenes*. Three aromatic compounds – toluene, benzaldehyde and ethylbenzene – also exhibited marked species-specific profiles. Ethylbenzene was present at trace levels only in *S. pyogenes* culture (Supplemental material Fig. 1). However, benzaldehyde levels were profoundly higher (24-fold) in *S. pyogenes* compared to *S. intermedius and S. pneumoniae* (Figure 4), suggesting a highly distinct metabolic route in *S. pyogenes*.

Any discriminating bacterial volatile biomarkers must be produced during the natural course of human infection or colonization. As all three bacterial species are natural colonizers of the upper respiratory tract, we hypothesized that volatiles produced by these bacteria should likewise be present in unbiased metabolomics of either human saliva or breath. Importantly, of the 26 discriminatory volatiles identified, we find that 11 of these (43%) have previously been characterized in metabolomics analyses of human saliva (22). While four have been proposed to be of dietary origin and previously reported in human breath (23) (2-nonanone; 2,3-pentanedione, 3-methylbutanal, and toluene), the remainder do not have a known human metabolic or exogenous origin. Our findings thus suggest that at least some component of the human salivary (and, by extension, human oropharyngeal) metabolomic fingerprint is likely to arise from the metabolic activities of the commensal oropharyngeal microbiota.

## DISCUSSION

Sore throat is one of the most common complaints encountered in the ambulatory clinical setting. Rapid, culture-independent diagnostic techniques that do not rely on pharyngeal swabs would be highly valuable as a point-of-care strategy to guide outpatient antibiotic treatment. Despite the promise of this approach, efforts to detect volatiles during oropharyngeal infection have yet been limited. Our study provides key proof-of-concept evidence that a unique pattern of volatile compounds is produced by *Streptococcus pyogenes*, the most common etiologic agent of bacterial pharyngitis. Importantly, we identify specific volatiles that distinguish this species from other typical human oropharyngeal colonizing bacteria. Our work thus lays the foundation for future clinical investigation into breath volatile detection as a mechanism to quickly and efficiently distinguish *S. pyogenes* pharyngeal infections.

For diagnostic purposes, the ideal volatile biomarker for *S. pyogenes* would be exclusively produced by this organism and only during conditions of active infection, rather than colonization. Because clinical studies of individuals with pharyngitis symptoms are both costly and time-consuming, it is important to build an understanding of the metabolic origin of bacterially produced volatiles, in order to predict whether candidate biomarkers will have the requisite sensitivity and specificity necessary for ongoing diagnostic development.

Linking experimentally determined metabolic capabilities back to bacterial genomes will ultimately build a predictive model for bacterial species that have yet to be characterized by volatile “fingerprint”. To date several studies have described the volatile patterns produced by model human pathogens (mostly Gram negative organisms) (7, 24), however, critical gaps exist in our understanding of the metabolic profiles of Gram positive organisms and human commensal bacteria. To our knowledge, only one such study has addressed *S. pyogenes*-specific volatiles (*25*). However, this targeted analysis only evaluated 10 volatile compounds that were present above the levels of quantification. In another study, the volatile headspace composition above Detroit cells inoculated with influenza A virus and *S. pyogenes* was investigated (26). Authors found significant differences in emitted VOC concentrations between non-infected and co-infected cells; however, the VOC profiling of *S. pyogenes* alone in cells was not tested. Similarly, the volatile profile of cultured *Streptococcus pneumoniae* has also been previously reported (19, 27), although these volatiles have yet to be directly compared to those from other oropharyngeal bacterial species. In this study, we take advantage of significant technological advancements that now provide a holistic and unbiased approach to identification of discriminant volatile biomarkers.

While the exact molecular mechanism of biosynthesis of many bacterial volatiles remains unclear, our study suggests that volatile production in streptococci may be particularly influenced by amino acid metabolism. Bacterially produced aromatic compounds, such as benzaldehyde, toluene, and ethylbenzene, are believed to be generated through enzymatic degradation of aromatic amino acids (e.g., phenylalanine, tryptophan, and tyrosine) (28). In addition, nitrogen-containing compounds represented the largest group of discriminant volatiles among the three streptococcal species under study. Several of these compounds (compounds 9, 10, 11, and 12; see Table 2) are aromatic heterocycles known as pyrazines. Pyrazines have strong odor properties, are widespread across phyla, and are among the most common classes of bacterially-produced volatiles (29). For this reason, pyrazines are of particular interest as biomarkers of bacterial infection. While pyrazines can be formed non-enzymatically (e.g., during autoclaving), the pattern of production of the discriminant pyrazines identified in our study (elevated in all three *S. pyogenes* isolates, but not *S. pneumoniae* or *S. intermedius*) strongly suggests an active biosynthetic role of *S. pyogenes*. Study of insect-associated *S. marcescens* revealed a bacterial enzymatic origin for pyrazine metabolites, with metabolic labeling confirming that these compounds derive from amino acid metabolism (specifically L-threonine) (30). More recently, the metabolic mechanism of pyrazine production in *Pseudomonas fluorescens* has been elucidated, confirming an enzymatic route that catalyzes the conversion of alpha amino acids to pyrazines (31). Our work provides strong evidence that *S. pyogenes*—but not *S. pneumoniae* or *S. intermedius*—possesses the molecular machinery for pyrazine biosynthesis. However, additional study is required to establish whether this enzymatic route is similar between *S. pyogenes* and *P. fluorescens*, as the first dedicated enzyme of the pyrazine biosynthesis pathway of *P. fluorescens*, PapD (PFLU_1773), does not have a close homolog in *S. pyogenes*.

Similarly, branch aldehyde production likely results from the catabolism of amino acids, a major energy source in TH media (32). Aldehydes may then be reduced to alcohols by alcohol dehydrogenases (e.g. 3-methylbutanal to 3-methyl-1-butanol) or oxidized to carboxylic acid by aldehyde dehydrogenase. Since 3-methylbutanal and 3-methyl-1-butanol were found to be released by all streptococcal species tested, our data suggest that amino acid degradation, rather than byproducts of fatty-acids (FA) metabolism, is responsible for the underlying pattern of VOCs released by these species. Microbially derived short-chain-branched alcohols, such as 3-methyl-1-butanol, are likely produced by enzymatic conversion of branched-chain amino acids (e.g., leucine and isoleucine) via the Ehrlich pathway (33).

In contrast, acetone is a ketone volatile that is present at high levels in all samples (second highest compound in concentration). While acetone is unsuitable as a biomarker due to its high concentrations in normal breath (34), the ketone 2,3-pentanedione has potential as a specific biomarker, and according to the Kegg pathway, *S. pyogenes* has the genetic capacity to produce it upon cysteine metabolism. Moreover, this ketone has been previously detected in the saliva of healthy individuals (35), and it is biologically plausible that streptococcal infection of the oropharynx could result in increased levels of 2,3-pentanedione.

To optimize the likelihood of identifying discriminant volatiles between streptococcal species, despite their metabolic similarities, we chose a defined growth condition to reduce the impact of contribution of differential growth rates to the differences in bacterially produced volatiles. As a result, the current study is limited by its investigation of this specific culturing condition, and additional study will be required to evaluate whether these volatiles are similarly discriminant during *in vivo* colonization or natural human infection. Furthermore, the current study is naturally limited to identification of volatiles arising from bacterial metabolism. During natural human infection, it is likely that host-derived volatiles will be present. These host-derived volatiles may be particularly important for diagnosis of *S. pyogenes*, as asymptomatic colonization is common (up to 20%) (36) and does not require treatment, unlike acute infection to which the host has a robust immune and tissue response.

In summary, our study provides evidence to pursue ongoing study of volatile biomarkers to discriminate oropharyngeal streptococcal species. Of note, several of the discriminant compounds, notably pyrazine compounds, are normally absent in healthy human exhaled breath (35) but can be found in the headspace of *Streptococcus pyogenes* cultures. That these compounds are typically absent suggests that a number of these candidate biomarkers are not typically produced by other members in the oropharyngeal microbial community. Such *S. pyogenes*-specific volatiles thus show particular promise as volatile biomarkers of acute streptococcal pharyngitis. Ongoing studies are required to validate our candidate biomarkers in patients with and without naturally acquired acute bacterial pharyngitis. Multiple factors, such as metabolite availability and host interaction with the bacteria through inflammatory responses will likely alter bacterial volatile production in the setting of acute infection. The ability to confirm the detection of pathogenic infection by *S. pyogenes* in the airways using breath analysis is a very exciting and non-invasive approach that will help guide immediate and appropriate antibiotic use upon infection confirmation.

## Acknowledgment

We thank Dr. S. Celeste Morley (Washington University) and Dr. Malcolm Winkler (Indiana University) for providing strains of *S. pneumoniae*. We thank Greg Storch (Washington University) for helpful comments. Research reported in this publication was supported by NIH/NIAID AI103280 (AOJ), AI123433 (AOJ), AI130584 (AOJ), AI144472 (AOJ), and the St. Louis Children’s Hospital Foundation. Dr. John is an Investigator in the Pathogenesis of Infectious Diseases of the Burroughs Welcome Fund. We also thank the NIH/NIGMS-supported Biomedical Mass Spectrometry (MS) Research Resource (P41GM103422).

## REFERENCES

1. Shulman ST, Bisno AL, Clegg HW. 2014. Clinical Practice Guideline for the Diagnosis and Management of Group A Streptococcal Pharyngitis: 2012 Update by the Infectious Diseases Society of America (vol 55, pg e86, 2012). Clinical Infectious Diseases 58:1496–1496.

2. Luo R, Sickler J, Vahidnia F, Lee YC, Frogner B, Thompson M. 2019. Diagnosis and Management of Group a Streptococcal Pharyngitis in the United States, 2011-2015. Bmc Infectious Diseases 19.

3. Kronman MP, Zhou C, Mangione-Smith R. 2014. Bacterial Prevalence and Antimicrobial Prescribing Trends for Acute Respiratory Tract Infections. Pediatrics 134:E956–E965.

4. Anonymous. 2017. CDC. Antibiotic Use in the United States, 2017: Progress and Opportunities. Atlanta, GA: US Department of Health and Human Services, CDC,

5. Hettinga KA, van Valenberg HJF, Lam TJGM, van Hooijdonk ACM. 2008. Detection of mastitis pathogens by analysis of volatile bacterial metabolites. Journal of Dairy Science 91:3834–3839.

6. Tait E, Perry JD, Stanforth SP, Dean JR. 2014. Use of volatile compounds as a diagnostic tool for the detection of pathogenic bacteria. Trac-Trends in Analytical Chemistry 53:117–125.

7. Bos LDJ, Sterk PJ, Schultz MJ. 2013. Volatile Metabolites of Pathogens: A Systematic Review. Plos Pathogens 9.

8. Bean HD, Dimandja JMD, Hill JE. 2012. Bacterial volatile discovery using solid phase microextraction and comprehensive two-dimensional gas chromatography-time-of-flight mass spectrometry. J Breath Res 901:41–46.

9. Berna AZ, McCarthy JS, Wang RX, Saliba KJ, Bravo FG, Cassells J, Padovan B, Trowell SC. 2015. Analysis of breath specimens for biomarkers of *Plasmodium falciparum* infection. J Infect Dis 212:1120–1128.

10. Schaber C, Katta N, Bollinger LB, Mwawi M, Mlotha-Mitole R, Trehan I, Raman B, Odom John A. 2018. Breathprinting reveals malaria-associated biomarkers and mosquito attractants. J Infect Dis 217:1553–1560.

11. Wilson AD, Baietto M. 2011. Advances in electronic-nose technologies developed for biomedical applications. Sensors 11:1105–1176.

12. Phillips M, Basa-Dalay V, Blais J, Bothamley G, Chaturvedi A, Modi KD, Pandya M, Natividad MPR, Patel U, Ramraje NN, Schmitt P, Udwadia ZF. 2012. Point-of-care breath test for biomarkers of active pulmonary tuberculosis. Tuberculosis 92:314–320.

13. Sreekumar J, France N, Taylor S, Matthews T, Turner P, Bliss P, Brook AH, Watson A. 2015. Diagnosis of Helicobacter pylori by carbon-13 urea breath test using a portable mass spectrometer. SAGE open medicine 3:2050312115569565–2050312115569565.

14. McFarland M, Szasz TP, Zhou JY, Motley K, Sivapalan JS, Isaacson-Schmid M, Todd EM, Hogan PG, Fritz SA, Burnham CAD, Hoffmann S, Morley SC. 2017. Colonization with 19F and other pneumococcal conjugate vaccine serotypes in children in St. Louis, Missouri, USA. Vaccine 35:4389–4395.

15. Wang XR, Cassells J, Berna AZ. 2018. Stability control for breath analysis using GC-MS. Journal of chromatography B 1097-1098:27–34.

16. Anonymous. 2008. NIST Standard Reference Database 1A, User Guide. The NIST Mass Spectrometry Data Center, National Institute of Standards and Technology,

17. Tautenhahn R, Patti GJ, Rinehart D, Siuzdak G. 2012. XCMS Online: A Web-Based Platform to Process Untargeted Metabolomic Data. Analytical Chemistry 84:5035–5039.

18. Metsalu T, Vilo J. 2015. ClustVis: a web tool for visualizing clustering of multivariate data using Principal Component Analysis and heatmap. Nucleic Acids Research 43:W566–W570.

19. Nizio KD, Perrault KA, Troobnikoff AN, Ueland M, Shoma S, Iredell JR, Middleton PG, Forbes SL. 2016. In vitro volatile organic compound profiling using GCxGC-TOFMS to differentiate bacteria associated with lung infections: a proof-of-concept study. Journal of Breath Research 10.

20. Phan J, Meinardi S, Barletta B, Blake DR, Whiteson K. 2017. Stable isotope profiles reveal active production of VOCs from human-associated microbes. Journal of Breath Research 11.

21. Hosmer J, McEwan AG, Kappler U. 2023. Bacterial acetate metabolism and its influence on human epithelia. Emerging Topics in Life Sciences doi:10.1042/etls20220092.

22. Zhang B, Hu S, Baskin E, Patt A, Siddiqui JK, Mathé EA. 2018. RaMP: A Comprehensive Relational Database of Metabolomics Pathways for Pathway Enrichment Analysis of Genes and Metabolites. Metabolites 8:16.

23. Amann A, Costello B, Miekisch W, Schubert J, Buszewski B, Pleil J, Ratcliffe N, Risby T. 2014. The human volatilome: volatile organic compounds (VOCs) in exhaled breath, skin emanations, urine, feces and saliva. Journal of Breath Research 8:034001.

24. Dolch ME, Hornuss C, Klocke C, Praun S, Villinger J, Denzer W, Schelling G, Schubert S. 2012. Volatile compound profiling for the identification of Gram-negative bacteria by ion-molecule reaction–mass spectrometry. Journal of Applied Microbiology 113:1097–1105.

25. Thorn RMS, Reynolds DM, Greenman J. 2011. Multivariate analysis of bacterial volatile compound profiles for discrimination between selected species and strains in vitro. Journal of Microbiological Methods 84:258–264.

26. Traxler S, Barkowsky G, Sass R, Klemenz AC, Patenge N, Kreikemeyer B, Schubert JK, Miekisch W. 2019. Volatile scents of influenza A and S. pyogenes (co-)infected cells. Scientific Reports 9.

27. Preti G, Thaler E, Hanson CW, Troy M, Eades J, Gelperin A. 2009. Volatile compounds characteristic of sinus-related bacteria and infected sinus mucus: Analysis by solid-phase microextraction and gas chromatography-mass spectrometry. Journal of Chromatography B-Analytical Technologies in the Biomedical and Life Sciences 877:2011–2018.

28. Carmona M, Zamarro MT, Blazquez B, Durante-Rodriguez G, Juarez JF, Valderrama JA, Barragan MJL, Garcia JL, Diaz E. 2009. Anaerobic Catabolism of Aromatic Compounds: a Genetic and Genomic View. Microbiology and Molecular Biology Reviews 73:71–+.

29. Dickschat JS, Wickel S, Bolten CJ, Nawrath T, Schulz S, Wittmann C. 2010. Pyrazine Biosynthesis in Corynebacterium glutamicum. European Journal of Organic Chemistry doi:10.1002/ejoc.201000155:2687-2695.

30. Silva EA, Ruzzini AC, Paludo CR, Nascimento FS, Currie CR, Clardy J, Pupo MT. 2018. Pyrazines from bacteria and ants: convergent chemistry within an ecological niche. Scientific Reports 8.

31. Masuo S, Tsuda Y, Namai T, Minakawa H, Shigemoto R, Takaya N. Enzymatic cascade in Pseudomonas for pyrazine production from α-amino acids. ChemBioChem 0.

32. Marilley L, Casey MG. 2004. Flavours of cheese products: metabolic pathways, analytical tools and identification of producing strains. International Journal of Food Microbiology 90:139–159.

33. Connor MR, Liao JC. 2008. Engineering of an Escherichia coli strain for the production of 3-methyl-1-butanol. Applied and Environmental Microbiology 74:5769–5775.

34. Schwarz K, Pizzini A, Arendacka B, Zerlauth K, Filipiak W, Schmid A, Dzien A, Neuner S, Lechleitner M, Scholl-Burgi S, Miekisch W, Schubert J, Unterkofler K, Witkovsky V, Gastl G, Amann A. 2009. Breath acetone-aspects of normal physiology related to age and gender as determined in a PTR-MS study. Journal of Breath Research 3.

35. de Lacy Costello BD, Amann A, Al-Kateb H, Flynn C, Filipiak W, Khalid T, Osborne D, Ratcliffe NM. 2014. A review of the volatiles from the healthy human body. Journal of Breath Research 8:1–29.

36. Roberts AL, Connolly KL, Kirse DJ, Evans AK, Poehling KA, Peters TR, Reid SD. 2012. Detection of group A Streptococcus in tonsils from pediatric patients reveals high rate of asymptomatic streptococcal carriage. BMC Pediatrics 12:3.

37. Lanie JA, Ng WL, Kazmierczak KM, Andrzejewski TM, Davidsen TM, Wayne KJ, Tettelin H, Glass JI, Winkler ME. 2007. Genome sequence of Avery’s virulent serotype 2 strain D39 of Streptococcus pneumoniae and comparison with that of unencapsulated laboratory strain R6. Journal of Bacteriology 189:38–51.

